# A Comparison of ‘Pruning’ During Multi-Step Planning in Depressed and Healthy Individuals

**DOI:** 10.1101/2020.05.08.084509

**Authors:** Paul Faulkner, Quentin J.M. Huys, Daniel Renz, Neir Eshel, Stephen Pilling, Peter Dayan, Jonathan P. Roiser

**Affiliations:** Department of Psychology, University of Roehampton, London, U.K.; Division of Psychiatry, University College London, London, U.K.; Max Planck Centre for Computational Psychiatry and Ageing Research, University College London, London, U.K.; Translational Neuromodeling Unit, Institute for Biomedical Engineering, University of Zurich and ETH Zurich, Zurich, Switzerland; Department of Psychiatry and Behavioral Sciences, Stanford University, Palo Alto, California, U.S.A; Division of Psychology and Language Sciences, University College London, London, U.K; Department of Computational Neuroscience, Max Planck Institute for Biological Cybernetics, Tübingen, Germany; Institute of Cognitive Neuroscience, University College London, London, U.K

## Abstract

**Background:** Real-life decisions are often complex because they involve making sequential choices that constrain future options. We have previously shown that to render such multi-step decisions manageable, people “prune” (i.e. selectively disregard) branches of decision trees that contain negative outcomes. We have theorized that sub-optimal pruning contributes to depression by promoting an oversampling of branches that result in unsavoury outcomes, which results in a negatively-biased valuation of the world. However, no study has tested this theory in depressed individuals.

**Methods:** Thirty unmedicated depressed and 31 healthy participants were administered a sequential reinforcement-based decision-making task to determine pruning behaviours, and completed measures of depression and anxiety. Computational, Bayesian and frequentist analyses examined group differences in task performance and relationships between pruning and depressive symptoms.

**Results:** Consistent with prior findings, participants robustly pruned branches of decision trees that began with large losses, regardless of the potential utility of those branches. However, there was no group difference in pruning behaviours. Further, there was no relationship between pruning and levels of depression/anxiety.

**Limitations:** The relatively small sample size limited the examination of individual differences. The use of other heuristics that are used to render complex decisions feasible, such as memoization and fragmentation, were not examined.

**Conclusions:** We found no evidence that sub-optimal pruning is evident in depression. Future research could determine whether maladaptive pruning behaviours are observable in specific sub-groups of depressed patients (e.g. in treatment-resistant individuals), or whether misuse of other heuristics may contribute to depression.

## Introduction

Major depressive disorder is a leading contributor to disability worldwide, affecting more than 300 million people at any time (World Health Organization, 2017). While negative affect and anhedonia are considered the predominant features of this disorder, people with depression also experience impairments in decision-making that may contribute to their depression (Clark et al., 2011; Eshel and Roiser, 2010; Husain and Roiser, 2018; Pulcu et al., 2015). Importantly, our knowledge of maladaptive decision-making processes in depression remains incomplete. Understanding such depression-related impairments may improve our comprehension of this disorder, and aid attempts to improve the quality of life for those that struggle with it.

Although people with anhedonic forms of depression are thought to exhibit dysfunctional evaluation of immediately-available, yet superficially appetitive outcomes, those with other forms of depression, such as those associated with helplessness, may exhibit maladaptive assessment of outcomes that become available only after more extended sequences of decisions (Dayan and Huys, 2008; Huys et al., 2015a). Addressing the substantial computational challenge associated with planning in multi-step decision-making problems lies at the heart of modern reinforcement learning (RL). In such cases it becomes necessary to search a decision-tree of options, in which the first choice constrains future choices. Such searches are difficult, and necessitate the use of certain heuristics.

Using the multi-step decision-making task in Figure 1, we have previously investigated various heuristics that healthy individuals use to approximate the best possible overall outcome without exhausting cognitive resources. One particular heuristic we identified is to ‘prune’ branches of a decision-tree that contain very negative outcomes in a reflexive (Pavlovian) manner (Huys et al., 2012; Huys et al., 2015b). Specifically, we found that healthy individuals are able to select the optimal sequence of decisions when potential paths do not contain a large loss, but are dramatically impaired at doing so when the optimal sequence involves incurring a large loss at the first decision step. In other words, healthy individuals prune branches that begin with a large loss, regardless of the potential net benefit of those branches (Figure 2).

**Figure 1.**
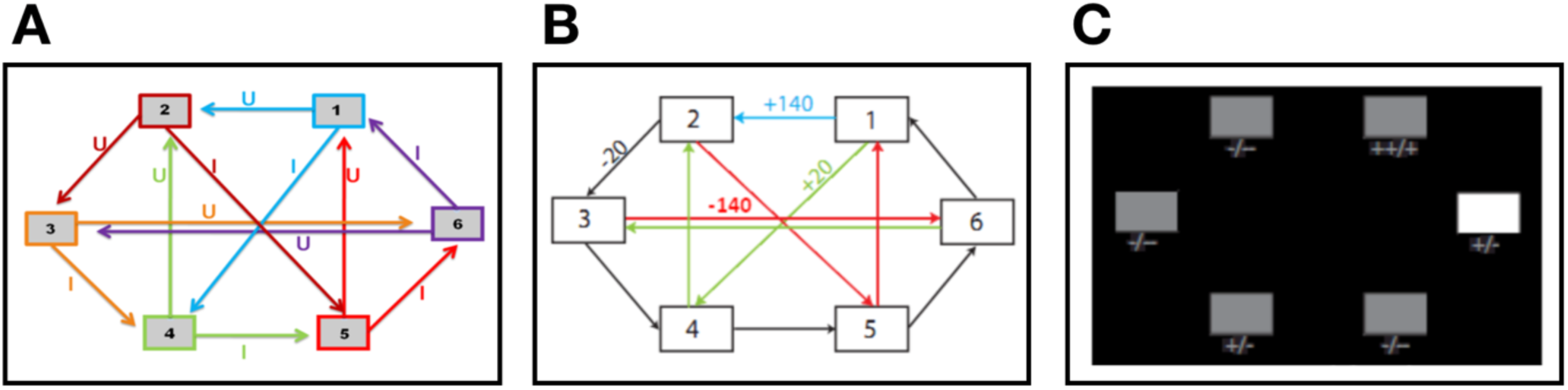
**A:** Deterministic transition matrix presented to participants during the training phase to aid learning. **B:** Deterministic reward matrix. Note that this was never presented to participants; they must instead learn the reward structure through trial and error. **C:** Task as presented to participants. The white box denotes the state that the participant is currently in. Symbols below each state denote the deterministic reward achieved by transitioning away from that state; ‘++’ = +140 points; ‘+’ = +20 points; ‘-’ = −20 points; ‘--’ = −140 points.

**Figure 2.**
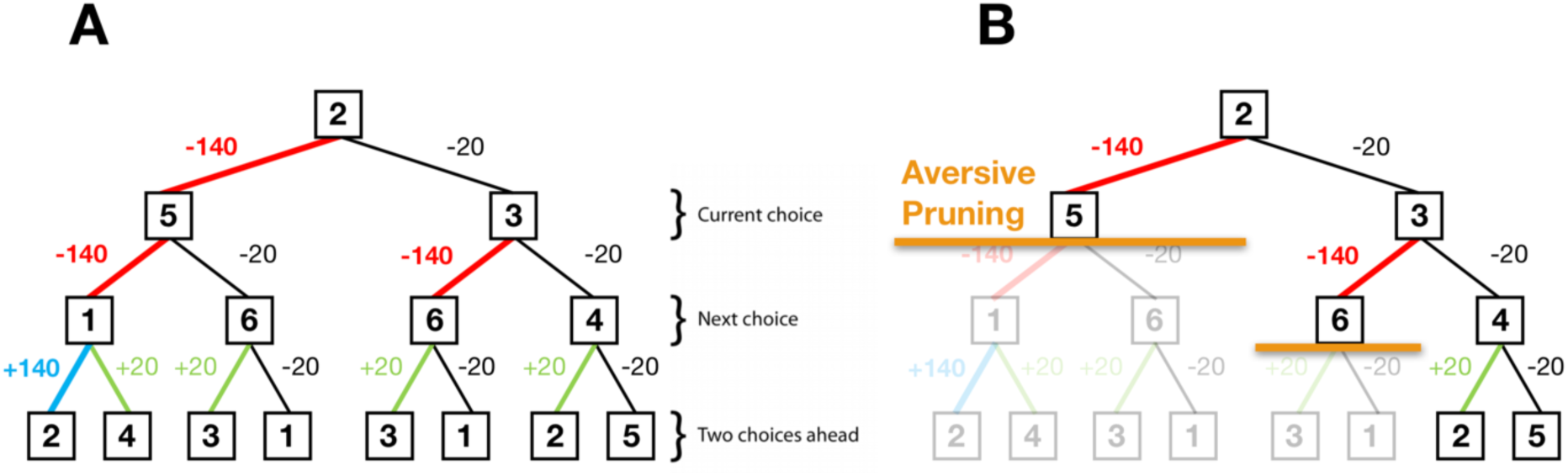
**A:** A typical decision tree and financial outcomes up to a depth of 3 starting from state 2. Numbers in each box denote the state number. **B:** Same decision tree starting, aversively pruned due to a large negative outcome at the first step. Note that this aversive pruning also avoids the large positive transition, but almost halves the computational load.

Pruning of decision branches that predict negative utility (i.e. branches that contain a large loss) can reflect a rational allocation of computational resources. However, such pruning also has a potential role in promoting emotional wellbeing; curtailing a tree search in the face of high negative utility avoids contemplation of possible aversive outcomes, which in turn may avoid a negatively-biased valuation of one’s surroundings (Dayan and Huys, 2008). Conversely, oversampling of branches that result in unsavoury outcomes may contribute to a more negatively-biased valuation of one’s surroundings and promote depression (Dayan and Huys, 2008; Huys et al., 2015b). In support of this hypothesis are findings that depression is associated with failures to inhibit processing of aversive information (Garrett et al., 2014; Joormann, 2005; Joormann and Gotlib, 2008). Indeed, in our prior work, individual differences in aversive pruning were related to the degree of sub-clinical depression in healthy participants (Huys et al., 2012).

Pavlovian pruning of decision trees has been hypothesized to depend on the brain’s serotonin system (Dayan and Huys, 2008). Specifically, reductions in central serotonin (whether naturalistic or experimental) are proposed to result in a decrease in Pavlovian behavioural inhibition, which usually acts to curtail internally-directed negative thoughts. Consequently, such decreases in serotonin are hypothesised to result in increased choosing of options that result in negative outcomes due to decreases in pruning (Dayan and Huys, 2008). This hypothesis was based on observations that serotonin influences behavioural inhibition in response to losses (Crockett et al., 2009; Cools et al., 2011). Because depression is associated with serotonergic dysfunction (Anderson et al., 2000; Cannon et al., 2007; Parsey et al., 2003), this theory provides one possible mechanism by which depressed individuals exhibit a negatively-biased valuation of themselves and their surroundings (Clifford and Hemsley, 1987).

The current study aimed to determine whether depression is associated with sub-optimal pruning of decision trees. We compared the performance of depressed and healthy participants on a sequential decision-making task designed to reveal pruning behaviours. Standard, Bayesian, and computational analyses of decision-making data were utilized. On the basis of previous work indicating a) that depression and serotonergic dysfunction are both associated with over-evaluation of negative information, and b) that depressed participants exhibit serotonergic hypofunction, it was predicted that 1) depressed participants would exhibit lower pruning of decision trees in the face of large punishments compared to healthy participants, and 2) that in depressed individuals, those with the highest levels of depression would demonstrate the lowest pruning.

## Methods

### Participants

This study employed a between-subjects design. Thirty depressed individuals were recruited via the Camden and Islington NHS Foundation Trust Psychological Therapy Services, while 31 healthy controls were recruited via online and print advertisements. All participants gave written informed consent after receiving a detailed explanation of the study (approved by the London - Queen Square NHS Research Ethics committee). Exclusion criteria (assessed by the Mini International Neuropsychiatric Inventory; MINI; Sheehan et al, 1998) for the healthy control participants included past or present major depressive disorder, bipolar disorder, psychosis, anxiety disorders, substance/alcohol dependence or recent (<6 months) abuse, any neurological disorder and not being a native English speaker. Depressed participants were subject to the same exclusion criteria, except they had to endorse depressive symptoms for a minimum of 10 days within the last two weeks (number of past depressive episodes was not important for inclusion), and could also have a diagnosis of an anxiety disorder, or historical substance/alcohol dependence that was restricted to a depressive episode. Exclusion criteria for depressed participants included use of antidepressants within the last month. All participants provided written informed consent and were compensated £30 for participation, as well as an additional £0-20 depending on their task performance.

### Procedure

Testing took place at the Institute of Cognitive Neuroscience, University College London. Participants completed one testing session each, in which they were administered the MINI and assigned to either the depressed or non-depressed group, depending on their responses. They then completed a battery of questionnaires (*see Questionnaire Measures, below*), before being trained on and performing a sequential decision-making task (*see Decision-Making Task, below*). Upon completion of the task, participants were paid according to their earnings.

### Questionnaire Measures

The Beck Depression Inventory (BDI; Beck, et al 1996) was administered to quantify severity of depression. This self-report questionnaire consists of 21-items, each of which is scored on a 0 - 3-point scale. A total score of 0-9 indicates minimal depression, while total scores of 10-18, 19-29 and 30-63 indicate ‘mild’, ‘moderate’ and ‘severe’ depression, respectively.

The State/Trait Anxiety Inventory (STAI; Speilberger et al., 1983) is a 40-item self-rating anxiety measure. The first 20 items quantify state anxiety, and the second 20 items quantify trait anxiety. Participants score each item as either 1 (‘do not agree at all’), 2 (‘agree somewhat’), 3 (‘agree moderately’) or 4 (‘very much agree’). Scores are then summed to provide separate total state and trait anxiety scores.

The Wechsler Test of Adult Reading (WTAR; Wechsler et al., 2001) was used to quantify verbal IQ. Participants were required to read aloud a list of 50 increasingly-uncommon words. A point was given for each correct pronunciation, and verbal IQ was calculated following conversion to standardised scores.

### Decision-Making Task

This task is described in Huys et al (2012), and is presented in *Figure 1*. Participants were required to complete both a training phase, and a task phase. During the training phase, participants were presented with a matrix containing 6 boxes (‘states’), and could make deterministic transitions between these states by pressing U or I on the keyboard. Participants learned the rules of the transitions by referring to the schematic of the transition matrix presented in *Figure 1a*, unlike subjects in Huys et al (2012) who learned these transitions by trial and error (i.e. without being presented with the schematic of the transition matrix). Importantly, as in Huys et al (2012), participants were only allowed to proceed to the task phase after demonstrating successful learning by completing a test without the aid of this schematic.

The task phase began with a further, shorter, training phase designed to teach participants the deterministic financial outcomes associated with each transition. Participants were not presented with a schematic of the action-reward matrix (see *Figure 1b*), but rather learned via trial and error. To help them, the potential values of the deterministic financial outcomes associated with each transition out of a state were depicted symbolically below each state (but without being identified with the choice options ‘U’ or ‘I’), both during this final training phase and throughout the entire task; ‘++’ denotes a £1.40 gain, ‘+’ denotes a 20 pence gain, ‘--’ denotes a £1.40 loss, and ‘-’ denotes a 20 pence loss. Upon completion of this final training, participants completed 48 trials of the task, each of varying length (2-8 moves). On each trial, participants began in a random state and were instructed to complete a sequence of transitions of a pre-specified length (2-8 moves) to maximize financial gain. On 50% of the trials, transitions were made immediately after each key press. followed by presentation of the relevant financial outcome for that transition in the centre of the screen. On the remaining trials (termed ‘plan-ahead’ trials), participants were instructed to plan ahead the remaining (2-4) moves and complete the full sequence of transitions, one immediately after another; the full sequence of transitions and resultant financial outcomes were only presented after the final key press had been made. Neither the transition nor reward matrices changed during the training or task phases. For clarity, a schematic of the decision tree when starting in state 2 can be seen in Figure 2a.

### Model-Based Statistical Analyses

To replicate the procedure in Huys et al (2012), the first 24 of the 48 trials were considered an extension of the reward matrix training, and data from these trials were therefore not analysed. A set of eight increasingly complex models was fit to the data using a Bayesian model comparison approach, all of which are fully defined in Huys et al (2012). Briefly, each successive model had an extra parameter to explain the data, and was assessed according to its Bayesian Information Criterion (BIC_int_), which is based on the likelihood function (i.e. the likelihood that the model can accurately explain the data) and penalizes the model for its extra complexity. A full description of these models can be observed in Huys et al (2012).

The first model was a simple ‘Lookahead’ model, which assumed that subjects evaluate the entire decision-tree to choose the optimal sequence of transitions, rather than pruning branches of the tree. That is for a trial of length *d*, this model assumed that participants considered all 2^*d*^ possible sequences of transitions and chose the financially most beneficial sequence. Because an explicit search all the way to the end of a tree is unlikely for depths >3 due to the large computational demands, more complex models were fit to the data, which allowed for testing of hypotheses pertaining to pruning.

The first of these, termed the ‘Discount’ model, built upon the previous model by including a ‘discount’ factor that assumed participants exhibit a general tendency to fail to evaluate all 2^*d*^ (i.e. up to 2^8^ = 256) sequences. Specifically, this model assumed that evaluation stopped at any stage along a path with probability γ. This is equivalent to the standard model of exponential discounting in economics, and implies that the deeper into the tree an outcome, the progressively less likely it is to be included in the calculation.

The next model, termed the ‘Pruning’ model, is central to this study’s hypotheses. This model splits the γ parameter from the previous model (i.e. the parameter that quantifies participants’ tendency to stop their tree search) into two separate parameters. The first of these is a ‘general pruning’ parameter (γ_G_) that quantifies the proportion of trials on which participants stopped their tree search due to a general tendency to fail to look ahead (identical to the ‘discount’ parameter in the previous model). The second is termed a ‘specific pruning’ parameter (γ_S_) that quantifies the proportion of trials on which the tree search was stopped specifically when the next transition would result in a large loss (see *Figure* 2b).

The next model, termed the ‘Pruning and Pavlovian’ model, accounted for ‘Learned Pavlovian’ attraction or repulsion to states that are associated with future financial rewards that may not be achievable on a particular trial (for example transitioning from state 6 to state 1 with one move remaining, despite not having enough moves remaining to achieve the +140 gain from subsequently transitioning out of state 1 to state 2). Specifically, it accounted for such learning due to the addition of a second state-action value which depends on the long-term average value of the states, which is itself learned by standard temporal difference learning after multiple exposures

Finally, to distinguish the effect of pruning from the effect of loss aversion (i.e. the notion that a loss of a given amount is more aversive than a reward of the same amount is appetitive), we replicated the above four models but relaxed how the models treated the different outcomes. In the original model, preferences for the outcomes were assumed to be proportional to the number displayed on the screen, and a loss of £1.40 was assumed to exactly cancel out a gain of £1.40. In the new models, we fitted separate parameters for each of the four financial outcomes, so that individuals could weight gains and losses differently, and indeed even weigh gains of different sizes in a non-proportional manner. These four models are termed ‘rho’ (i.e. ‘Lookahead rho’, ‘Discount rho’, ‘Pruning rho’ and ‘Pruning & Pavlovian rho’), with each having three additional parameters (a parameter for each outcome, but no overall scaling parameter as in the original models). In principle, this allowed participants to be attracted to a reward and repelled from a loss, and vice versa. If pruning is observable above and beyond an individual’s simple preferences for rewards and losses, the differential sensitivities to rewards and punishments cannot, by themselves, account for the pruning effects in the above four (i.e. non-’rho’) models.

### Group Comparisons and Psychometric Correlation Analyses

Once the best fitting model was identified, its parameter estimates were extracted and compared between the groups. These analyses were performed using both frequentist and Bayesian approaches. Frequentist analyses were performed using the Statistical Package for Social Scientists version 26 (SPSS Inc., Chicago, Illinois). Bayesian analyses were performed using JASP (JASP Team (2019), version 0.11.1). Bayesian analyses were performed because they provide Bayes Factors, which depict a ratio of the probability of the evidence for one hypothesis (i.e. the null) relative to another (i.e. the experimental). Comparing evidence in this way allows one to demonstrate support for the null hypothesis, as opposed to simply failing to reject the null as when using frequentist approaches (Wetzels et al., 2011).

To determine the effects of depression status on parameter estimates from the most parsimonious model, frequentist and Bayesian independent samples t-tests were performed, with the relevant parameter estimate added as the dependent variable. To examine relationships between parameter estimates and psychometric questionnaire data, frequentist and Bayesian bivariate correlation analyses were performed.

For the frequentist analyses, a significance threshold of alpha = 0.05 (2-tailed) was adopted. Bonferroni correction (BC; for 6 parameter estimates*3 questionnaire variables) was applied to the correlational analyses. For the Bayesian analyses, on the basis of Jeffreys (1961), we considered Bayes Factors (*BF*_10_; NB: not logarithmically transformed) smaller than 1/100 to be extreme evidence for the null hypothesis, a *BF*_10_ between 1/100 and 1/30 to be very strong evidence for the null, a *BF*_10_ between 1/30 and 1/10 to be strong evidence for the null, a *BF*_10_ between 1/10 and 1/3 to be moderate evidence for the null, and a *BF*_10_ between 1/3 and 1 to be not worth more than a bare mention. Conversely, we considered *BF*_10_ larger than 100 to be extreme evidence for the experimental hypothesis, a *BF*_10_ between 100 and 30 to be very strong evidence for the experimental hypothesis, a *BF*_10_ between 30 and 10 to be strong evidence for the experimental hypothesis, a *BF*_10_ between 10 and 3 to be moderate evidence for the experimental hypothesis, and a *BF*_10_ between 3 and 1 to be not worth more than a bare mention.

## Results

### Participant Characteristics

Compared to healthy controls, depressed participants self-reported higher depression scores on the BDI (t(59)=13.779; *p*<0.001, *BF*_10_=1.11e+16), greater trait anxiety on the STAI (t(59)=-7.601; *p*<0.001, *BF*_10_=1.672e+7), and greater state anxiety on the STAI (t(59)=-5.219; *p*<0.001, *BF*_10_=4987.46). Frequentist t-tests failed to reject the null hypothesis that depressed and healthy individuals did not differ in terms of age, years of education or IQ, although Bayesian analyses failed to provide support for or against the null hypothesis that these two groups did not differ in these characteristics; (age: t(59)=1.227; *p*=0.225, *BF*_10_=0.496; years of education: t(59)=1.491; *p*=0.141, *BF*_10_=0.660; IQ: t(59)=1.780; *p*=0.083, *BF*_10_=1.058). A full description of participant characteristics is presented in Table 1.

**Table 1.** Participant Characteristics. ^α^ Denotes mean (SD)

|  | Healthy<br>Individuals | Depressed<br>Individuals |
| --- | --- | --- |
| *N* | 31 | 30 |
| Gender (*M/F*) | 15/16 | 15/15 |
| Age (years)<sup>α</sup> | 30.42 (10.80) | 33.82 (10.44) |
| Education (years) <sup>α</sup> | 14.38 (2.94) | 15.43 (2.51) |
| IQ <sup>α</sup> | 108.00 (14.79) | 114.55 (7.37) |
| BDI | 3.00 (4.61) | 25.44 (7.48) |
| Trait Anxiety | 14.71 (8.16) | 32.88 (9.74) |
| State Anxiety | 10.19 (7.53) | 23.44 (11.39) |
| No. of depressive episodes<sup>α</sup> | - | 8.10 (3.82) |
| Current episode length (months) <sup>α</sup> | - | 3.64 (4.91) |
| No. of days depressed within past 2 weeks<sup>α</sup> | - | 10.63 (2.86) |
| Number of suicide attempts<sup>α</sup> | - | 0.55 (1.21) |

**Table 2.**
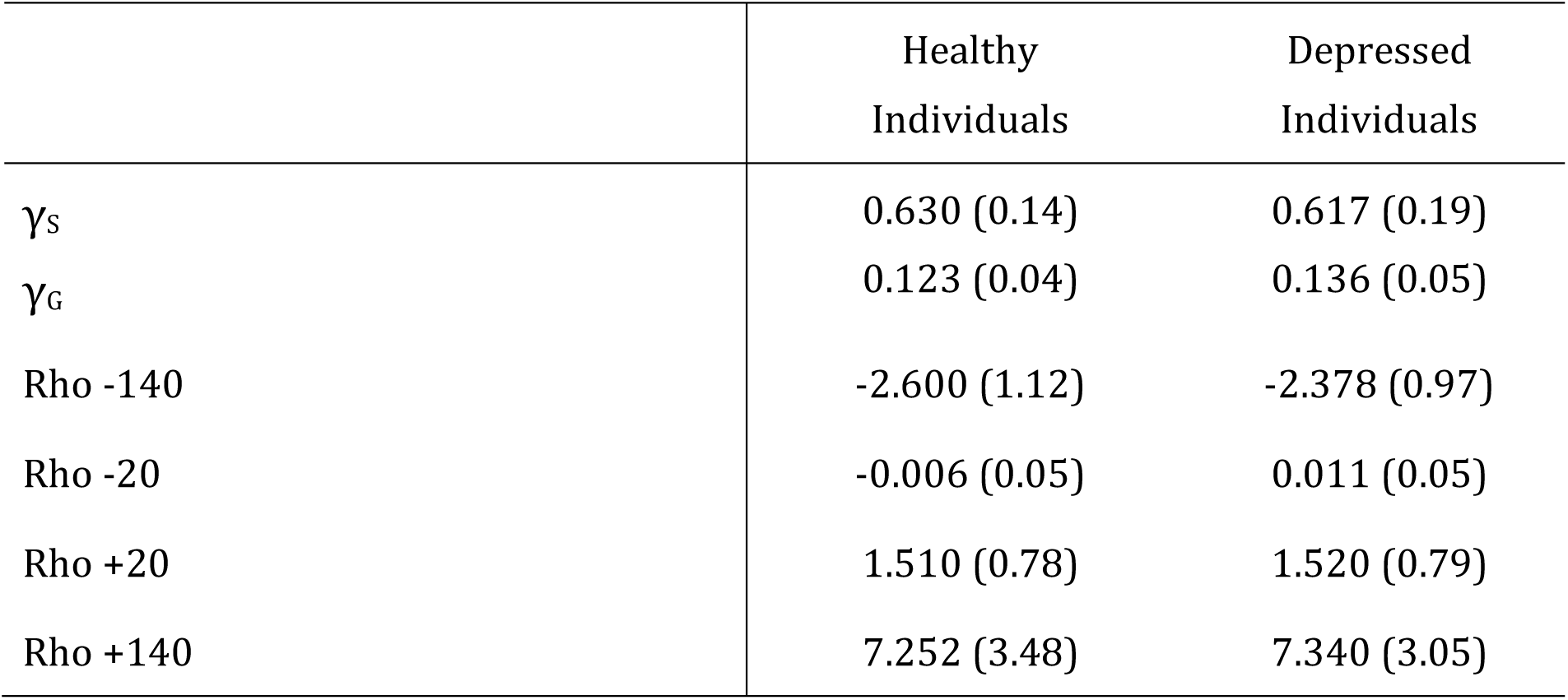
Parameter Estimates from the Winning Pruning ‘rho’ Model. All values are expressed as means (SD)

### Computational Analyses

The ability of all eight models to explain participants’ choices can be seen in *Figure* 3. The inclusion of each extra parameter improved the predictive probabilities of the models, while the four models that incorporated the ‘rho’ parameter were all able to predict a higher proportion of participants’ choices than the corresponding models that did not include this parameter (*Figure 3a*). Importantly, the Pruning ‘rho’ model outperformed all others due to it achieving the lowest BIC_int_ score (*Figure 3b*). As expected, the most complex model, the Pruning and Pavlovian ‘rho’ model, achieves the highest predictive probability (i.e. it is able to accurately predict the highest proportion of participants’ choices, as shown in *Figure 3a*). However, it is penalised for its added complexity (*Figure 3b*). This means that the Pruning ‘rho’ model is considered the most parsimonious, and therefore the winning, model.

**Figure 3.**
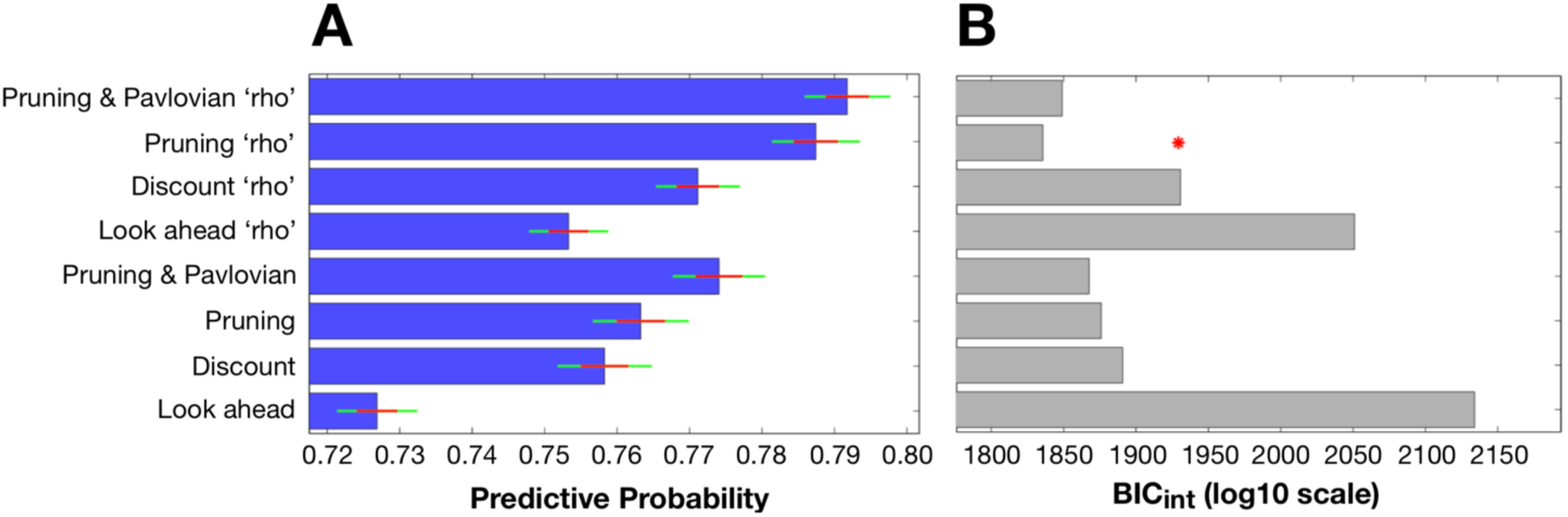
**A:** Mean predictive probabilities for all models. All models that include the ‘rho’ parameter fit the data better than the corresponding models that do not contain this parameter. **B:** Model comparisons using each model’s Bayesian Information Criterion (BIC_int_). Despite the fact that the model that predicts the highest proportion of participants’ choices is the ‘Pruning and Pavlovian’ model that contains the ‘rho’ parameter, this model is penalized due to its added complexity. The most parsimonious (i.e. ‘winning’) model is therefore the ‘Pruning’ model that includes the extra ‘rho’ parameter.

Specifically, the winning Pruning ‘rho’ model included both γ_S_ and γ_G_ parameters, which suggests that loss-specific pruning had a robust influence on behaviour. The fraction of choices that was correctly predicted by this winning model can be seen in *Figure 4a-c*. Because this Pruning ‘rho’ model outperforms all others, the γ_S_ and γ_G_ parameter values, as well as the reward sensitivities to each of the four transition types, were extracted from this model and compared between depressed and healthy participants.

**Figure 4.**
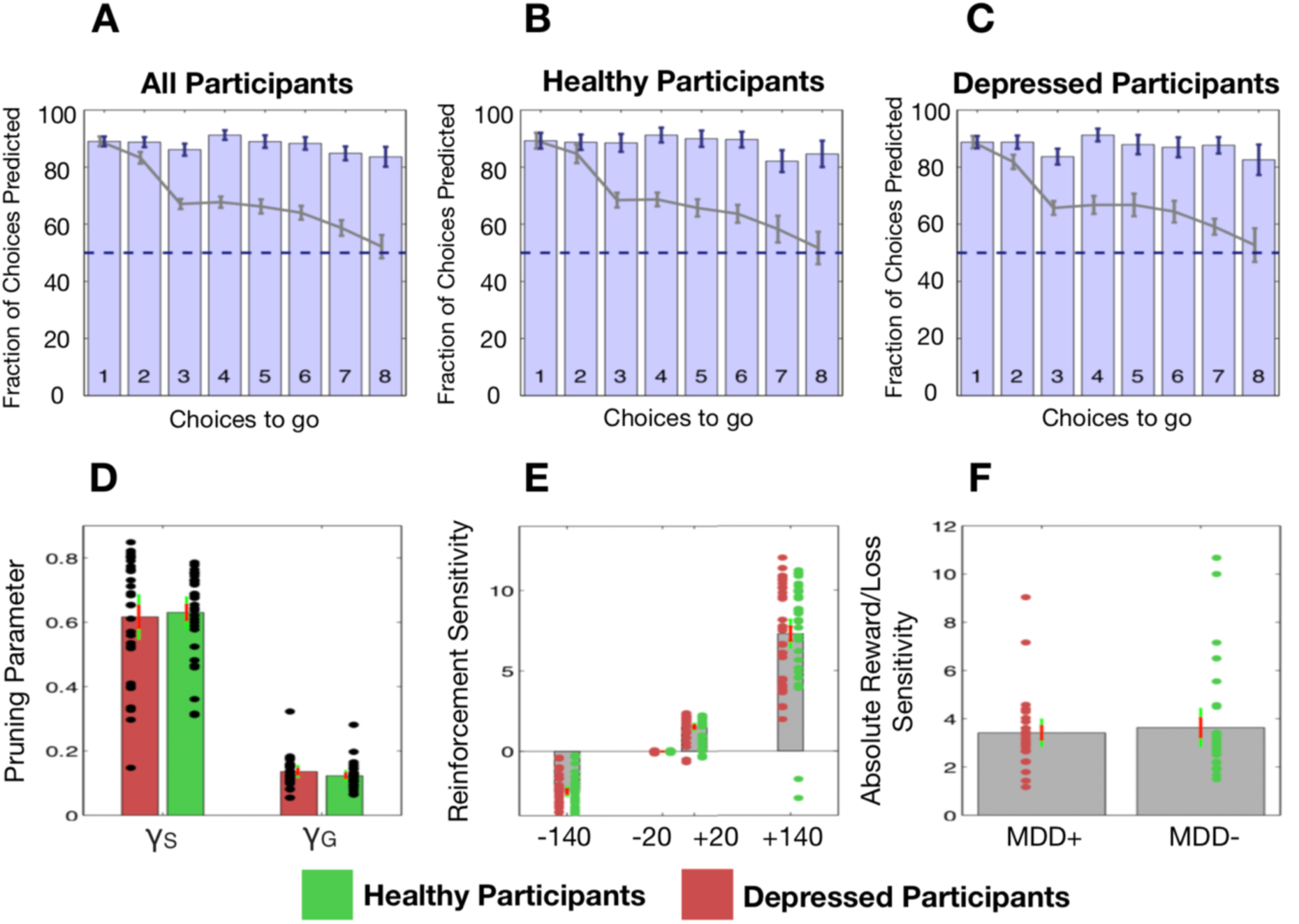
***Top:*** The fraction of choices correctly predicted by the best-fitting model (the Pruning ‘rho’ model). **A:** All participants combined. **B:** Healthy participants only. **C:** Depressed participants only. Each bar depicts this as a function of the number of choices remaining on each trial. For example, the right most bar (i.e. bar ‘8’) depicts the fraction of choices at a depth of 1 on 8-choice trials that were correctly predicted by this model; the third rightmost bar (i.e. bar ‘6’) depicts both the fraction of choices that were correctly predicted by this model at (1) a depth of 1 on 6-choice trials, (2) a depth of 2 on 7-choice trials and (3) a depth of 3 on 8-choice trials, and so on. Grey lines depict the full ‘Lookahead’ model. The blue dashed lines depict chance (i.e. 50%). The winning model correctly predicts choices of both depressed participants and healthy controls to roughly the same extent. Further, the full Lookahead model is only able to correctly predict decisions that are 8 choices away in the sequence on roughly 50% of trials (i.e. at chance level). The winning model correctly predicts all choices to roughly the same extent, no matter how many choices are remaining. Note that these models include data from, and disregard differences between, trials in which transitions were displayed immediately after each button press and trials in which participants had to enter the entire sequence of transitions at once (i.e. so-called ‘plan-ahead’ trials.). ***Bottom:*** Parameters of the winning Pruning ‘rho’ model. **D:** Specific and general pruning parameters. **E:** Reinforcement sensitivity to each transition type. **F:** Absolute ratio of reward (+140) to loss (−140) sensitivity. Red denotes depressed participants, green denotes healthy participants. Error bars denote 1 standard deviation above/below the mean (red) and 95% confidence intervals (green).

### Group Comparisons

Frequentist and Bayesian analyses indicated that depressed and healthy individuals did not differ in terms of total money won on the task (t(59)=0.345; *p*=0.731, BF_10_=0.274). Further, depressed and healthy individuals did not differ in terms of γ_S_ (t(59)=0.320; *p*=0.750, *BF*_10_=0.272,). However, while the frequentist analysis indicated that depressed and healthy individuals did not differ in terms of γ_G_ either, the Bayesian analysis failed to provide support for or against there being a group difference in this variable: (t(59)=-1.123; *p*=0.266, *BF*_10_=0.442,). These data can be seen in *Figure 4d*. There were no differences between the groups in terms of reward sensitivities for the +140 transitions (t(59)=-0.105; *p*=0.917, *BF*_10_=0.262), +20 transitions (t(59)=-0.073; *p*=0.942, *BF*_10_=0.261) or the −20 transitions (t(59)=0.459; *p*=0.648, *BF*_10_=0.285). Again, while a frequentist analysis indicated that depressed and healthy individuals did not differ in their sensitivity to the −140 transitions, the results of the Bayesian analysis narrowly missed out on providing support for the null hypothesis that these two groups did not differ in this variable (t(59)=-0.831; *p*=0.409, *BF*_10_=0.348). These data can be seen in *Figure 4e*.

In depressed participants, BDI scores did not correlate with γ_S_ (*r*=0.068; *p*=0.748, *BF*_10_=0.261), γ_G_ (*r*=-0.083; *p*=0.693, *BF*_10_=0.267), or reward sensitivities to the +140 transitions (*r*=0.038; *p*=0.857, *BF*_10_=0.252) or the −140 transitions (*r*=-0.137; *p*=0.514, *BF*_10_=0.304). Further, frequentist analyses indicated that BDI scores also did not correlate with the +20 transitions or −20 transitions, although the Bayesian analyses did not provide support for or against the null hypothesis that BDI scores did not correlate with the sensitivity to these two transition types (+20 transitions: *r*=-0.226; *p*=0.276, *BF*_10_=0.435; −20 transitions: *r*=0.261; *p*=0.207, *BF*_10_=0.528).

In addition, in depressed participants, trait anxiety scores did not correlate with γ_S_ (*r*=0.083; *p*=0.694, *BF*_10_=0.267), γ_G_ (*r*=0.084; *p*=0.688, *BF*_10_=0.268), or reward sensitivities to the +140 transitions (*r*=0.156; *p*=0.457, *BF*_10_=0.323), the +20 transitions (*r*=0.309; *p*=0.133, *BF*_10_=0.724), the −20 transitions (*r*=0.066; *p*=0.755, *BF*_10_=0.260) or - 140 transitions (*r*=-0.070; *p*=0.740, *BF*_10_=0.261).

Finally, both frequentist and Bayesian analyses indicated that state anxiety scores did not correlate with γ_S_ (*r*=-0.092; *p*=0.662, *BF*_10_=0.272) in depressed participants. However, while the frequentist analysis indicated that state anxiety did not correlate with γ_G_, results of the Bayesian analysis narrowly missed out on providing support for there being no relationship between these two variables (*r*=-0.177; *p*=0.397, *BF*_10_=0.349). Both frequentist and Bayesian analyses indicated that state anxiety was not related to sensitivities to the +140 transitions (*r*=-0.048; *p*=0.820, *BF*_10_=0.254), +20 transitions (*r*=-0.013; *p*=0.950, *BF*_10_=0.249), −20 transitions (*r*=-0.004; *BF*_10_=0.248, *p*=0.985) or −140 transitions (*r*=0.068; *p*=0.745, *BF*_10_=0.261).

## Discussion

We have previously shown that healthy individuals prune decision trees to render complex sequential decisions manageable. Here, we tested the hypothesis that depressed individuals are less able to optimally use this heuristic, and that this inability may be related to their severity of depression. However, contrary to predictions, the depressed participants and healthy individuals in this study did not differ in their pruning behaviours.

The finding that participants pruned branches of decision trees that began with a large loss, regardless of the potential utility of that branch, replicates our previous work (Huys et al., 2012; Huys et al 2015a). Further, the most parsimonious model in our original study (Huys et al., 2012) included a Pavlovian parameter which indicated that participants had a reflexive attraction/aversion to certain states. However, in the current study, including this Pavlovian parameter weakened model parsimony, meaning that participants did not display strong attractions/aversions to specific states in this dataset.

The addition of four separate parameters to the model (one for each of the four financial outcomes) allowed for the possibility that there was a difference between how participants weighted gains and how they weighted losses. This is important to avoid confusing pruning with loss aversion. Replicating the findings of Huys et al (2012), we found that these parameters (labelled as the ‘rho’ components) indeed improved the predictive probability of the models. However, as for the healthy individuals in Huys et al (2012), the current participants did not exhibit typical loss aversion in their decision-making. Instead, the large gain (+140) was roughly 3 times more appetitive than the large loss (−140) was aversive. Interestingly, sensitivity to −140 transition types was found to be significantly weaker, while sensitivity to the +140 transitions was significantly greater, in the ‘winning’ (Pruning ‘rho’) model than in the model that did not quantify participants’ pruning behaviours (i.e. the ‘lookahead’ model; see Supplementary Materials). Taken together, these above results suggest that loss aversion certainly fails to explain away pruning, although pruning may uncover a form of risk seeking.

Importantly, the current findings failed to support our hypotheses that a) depressed individuals sub-optimally prune decision trees, and that b) in depressed individuals, those with the highest levels of depression would demonstrate the lowest pruning. Our latter hypothesis was based partly on the results of Huys et al (2012), which indicated a relationship between pruning and level of depression. However, it must be noted that the specific relationship reported in Huys et al (2012) was a significant *positive* correlation between specific pruning and *sub-clinical* depression scores on the BDI in *healthy* participants, not depressed individuals. Further, this relationship was also not replicated by our subsequent studies (i.e. Huys et al 2015a). While it (so far) appears that no relationship between pruning and magnitude of depression exists in depressed individuals, future research should attempt to determine the replicability of the initial finding of a relationship between specific pruning and sub-clinical depression in healthy individuals. Further, regarding the fact that there was no group difference in pruning, it is unlikely that we failed to observe sub-optimal pruning in the current depressed individuals because the current healthy participants also pruned sub-optimally, because the latter demonstrated specific pruning to a very similar magnitude as the healthy participants included in Lally et al (2017) (pruning parameter estimate = ∼0.6), and they actually pruned slightly more than those in Huys et al (2012). While this does not explain why the current results do not support the theory put forward by Dayan and Huys (2008), there may be a number of factors that do.

First, levels of depression in our participants may not have been sufficiently high so as to promote sub-optimal pruning. While the current depressed participants exhibited similar levels of depression to those in studies that report maladaptive decision-making in depression (Joormann and Gotlib, 2008; Kumar et al., 2018; McFarland and Klein, 2009; Ubl et al., 2015), they were all undergoing ‘low intensity treatments’, including cognitive behavioural therapy, guided self-help and online support due to being deemed by psychologists to exhibit mild/intermediate levels of depression. Indeed, our depressed patients failed to exhibit greater sensitivity to losses than healthy participants, or indeed loss aversion at all. Highly depressed patients (Baek et al., 2017; with mean BDI = 30.10) have been shown to have heightened aversion to losses, unlike patients with lower levels of depression (Charpentier et al. 2017; mean BDI = 16.96). Future studies could therefore beneficially examine the pruning behaviours of more severely depressed participants.

Second, it was predicted that depressed participants would exhibit maladaptive pruning partly because optimal pruning is theorized to depend on serotonin, and because depression is associated with serotonergic hypofunction (Anderson et al., 2000). However, there is no way of knowing whether the current depressed participants exhibit such serotonergic dysfunction because no neurochemical measures were collected in this study. Further, depression has been associated with altered functioning of the brain’s dopamine system (Nestler and Carlezon, 2006), and this neurotransmitter is thought to influence decision-making in an opponent fashion to serotonin (Daw et al., 2002; Boureau and Dayan, 2010; Dayan and Huys, 2009). For example, experimentally-induced reductions in serotonin via acute tryptophan depletion can enhance the motivational influence of aversive stimuli on instrumental responding, while reductions in dopamine via acute tyrosine depletion can diminish the influence of appetitive stimuli on such responding (Hebart and Gläscher, 2014). To truly determine the influence of serotonergic function on pruning of decision trees, future studies may consider examining the effects of a serotonin challenge such as acute tryptophan depletion on performance on this task.

Third, while no group difference was observed in task performance, this does not mean that the brain regions that are recruited during aversive pruning show the same patterns of activity in depressed as healthy individuals. We have recently shown that aversive pruning recruits the pregenual anterior cingulate cortex (pgACC) and subgenual anterior cingulate cortex (sgACC) (Lally et al., 2017). Interestingly, the sgACC is considered to be overactive in depression (Drevets et al., 1997; 2008; Mayberg et al., 1997), while the degree of sgACC reactivity to negatively-valenced stimuli can predict treatment response in depressed patients (Roiser et al., 2012). In addition, depression-related reductions in serotonin 1A receptor availability are greatest in the sgACC (Moses-Kolko et al., 2008). On the basis of these findings, comparing the neural mechanisms of pruning in depressed and healthy individuals may reveal differences in pruning-related sgACC function between the two groups.

While depressed and healthy individuals did not differ in pruning, this is not the only heuristic people use to render complex, sequential decisions manageable. We have previously shown that healthy participants use two heuristics in addition to pruning. Specifically, participants solve problems by fragmenting deep sequences (i.e. sequences with a depth >3) into sub-sequences of shorter lengths (termed ‘fragmenting’), and recall and re-use previous fragmented solutions on subsequent trials (termed ‘memoization’), rather than always searching the tree anew each time (Huys et al., 2015a). Unlike the case for pruning (Dayan and Huys, 2008; Huys et al., 2012), neither theoretical nor empirical studies have suggested that altered use of fragmenting or memoization during planning may promote depression. However, future studies might compare the use of these two heuristics, along with the use of pruning, in highly-depressed and healthy individuals.

Our study has several limitations. First, the sample size was relatively limited, which reduced our ability to detect small effects with either Bayesian or frequentist methods. In addition, this relatively small sample size hindered our ability to determine the effects of individual differences on task performance; this could be particularly pertinent as Huys et al (2015a) report that individuals use certain heuristics to solve planning problems in an idiosyncratic fashion. Another potential limitation is the fact that our healthy and depressed individuals may have differed in age, years of education and I.Q. While frequentist analyses found no evidence that these two groups differed in such characteristics, Bayesian analyses provided, at best, only anecdotal evidence that these two groups did *not* differ in this way. As such, it is possible, although unlikely, that our sampling method may have introduced extraneous variables into our dataset that influenced our findings. Further, while we have reported that pruning is insensitive to the magnitude of the large loss (Huys et al., 2012)., pruning becomes mathematically more disadvantageous as the magnitude of the large loss decreases. However, the current data do not indicate whether depressed individuals prune sup-optimally when pruning is more disadvantageous (i.e. when the large loss costs 70 points rather than 140). Finally, our calculated reward sensitivities in figure 4E indicate that, once pruning is taken into account, participants are 3 times more ‘sensitive’ to the large reward than they were to the large loss. However, pruning and risk seeking are rather entangled in the current task, and it would be interesting to combine it with a compatible, but independent measure of risk seeking.

In summary, we replicated previous findings that people prune branches of decision trees to solve complex planning problems. However, we failed to provide support for the hypothesis that depressed individuals prune such branches sup-optimally. Future research is needed to achieve a more complete understanding of whether misuse of certain heuristics in sequential decision-making can contribute to depression.

## Supporting information

Supplementary Results

