## Supplementary Results for "A Comparison of ‘Pruning’ During Multi-Step Planning in Depressed and Healthy Individuals"

**Supplementary Materials**

There was a strong and meaningful difference between the reward sensitivities of the Pruning rho (i.e. 'winning') model and the Lookahead rho (i.e. baseline) model, in terms of the -140 transitions ($t(60) = 9.019$, $p < 0.001$, BF = 9.026e +9) and the +140 transitions ($t(60) = 12.317$, $p < 0.001$, BF = 1.245e +15), as well as in terms of the +20 transitions ($t(60) = 9.000$, $p < 0.001$, BF = 8.398e +9) but not the -20 transitions ($t(60) = 1.290$, $p = 0.202$, BF = 0.308). Specifically, compared to the Lookahead 'rho' model, participants sensitivities to the -140 and +140 transitions appear weaker and stronger, respectively, when pruning is taken into account (see Figure S1).

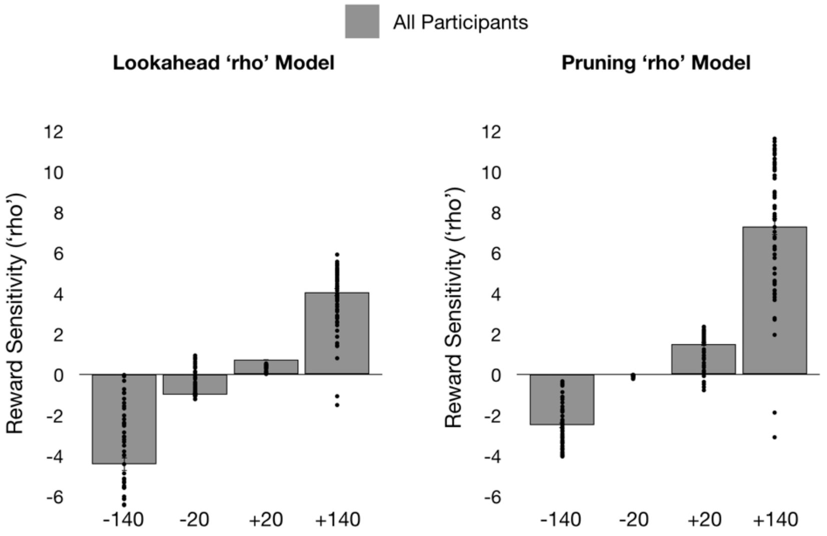


**Figure S1. Rho values for each of the four transition types in both the Lookahead 'rho' model (left) and the Pruning 'rho' model (right)**
